# In Rift Valley settings with a feedback loop, assortative mating for versatility predicts hominin brain enlargement in some detail

**DOI:** 10.1101/164780

**Authors:** William H. Calvin

**Affiliations:** Department of Psychiatry and Behavioral Sciences, University of Washington, 725 9^th^ Avenue, Box 2605, Seattle WA 98104 USA; | http://faculty.washington.edu/wcalvin; Center for Academic Research and Training in Anthropogeny, UCSD and the Salk Institute, La Jolla CA USA

**Keywords:** assortative mating, brain size, cerebral cortex, demography, habitat preference, hitchhiking, Homo erectus, language development, Neanderthal, neocortex, non-random gene flow

## Abstract

Hominin procedures for fire-starting, sharpening rocks, and softening roots by pounding or chopping require sustained attention for hours; shade is sought in the brush fringe bordering a grassland. Clustering these more versatile adults, while others are away hunting and gathering, provides a setup for assortative mating. This can lengthen attention span, enhance versatility and, with it, brain size. The rate of enlargement is accelerated by a boom-and-bust cycle in their meat supply, predicting the observed initiation of enlargement at −2.3 myr in the Rift Valley once boom-prone grazers evolved from the mixed feeders. Several months after lightning created a burn scar back in the brush, the new grassland enables a population boom for those grazers that discover it. Several decades later as brush regrows, they are pushed back. Their hominin followers, wicked in from the grassland’s shady fringe, boom together with the burn-scar grazers. They then follow their meat supply back to the main population. This creates an amplifying feedback loop, shifting *Homo* gene frequencies centrally. Brush fires are so frequent that the cosmic ray mutation rate becomes enlargement’s rate-limiter, consistent with 460 cm^3^/myr remaining constant during many climate shifts. The apparent tripling of enlargement rate in the last 0.2 myr vanished when the non-ancestors were omitted. Asian *Homo erectus* enlargement lags the ancestral trend line by 0.5 myr. Neanderthals lag somewhat less but have a late size spurt after the −70 kyr *Homo sapiens* Out of Africa, suggesting enlargement genes were acquired via interbreeding.

## 1. The other mechanism of natural selection

Evolution seems to have all of the permanence of a Heraclitan universe, though its theorists have attempted to fit economic principles to its froth and flow, hoping that efficiency arguments might illuminate the process because the inefficient die sooner. They tend to adopt the economists’ “There is no such thing as a free lunch.” But free rides exist and they are not mere analogues to the economists’ externalities.

### 1.1. Founder effect and its amplification

Natural selection does not operate solely by selective survival of useful mutations; there is an escape clause that has been obvious since the early days of evolutionary theory. Natural selection can also operate via selective reproductive success.

Consider the founder effect: whatever genes helped the parent to be in the right place at the right time for a range expansion promotes more surviving grandchildren carrying those genes. A habitat preference for living on the frontier, and occasionally being favored with more grandchildren than when in the core population, serves to increase the overall proportion interested in living on the frontier.

There are physiological versions as well: in those sheep variants where twinning rate temporarily increases with diet improvements (Heape 1899), as at the end of a drought, double ovulation serves to quadruple the number of grandchildren. This can easily offset a higher mortality rate when living on the frontier. Twins are risky when food is tight as both may be lost in conditions where singletons can survive.

Should the expanded population be squeezed back into the core population, it creates amplifying feedback (Fig. 1), opening up additional options for evolution. And because boom-and-bust are paired, one can see founder effects that repeat in the manner of the compounding of interest (Calvin 2017), making the nonrandom gene flow something closer to a favored-gene pump.

*Figure 1: Expand-squeeze gene pumping from boom-and-bust*

**Fig. 1.**
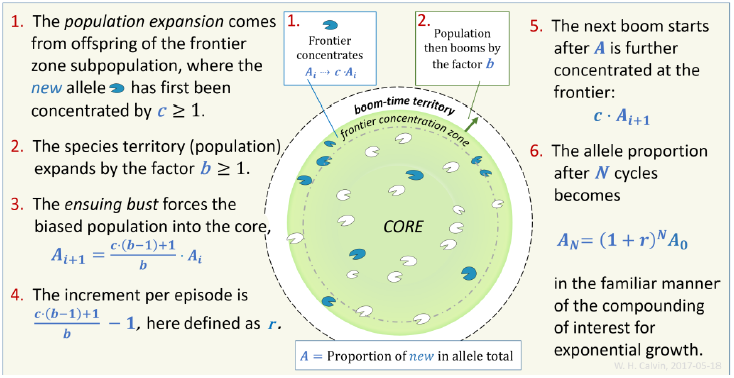
Expand-squeeze pumping illustrated with a simple center-surround diagram. The exponential compounding of allele proportions ends when ***A***_***N***_ = **1** (fixation). The constants *b* and *c* remain useful in other geometries such as the brush-fire burn scar. After Calvin (2017).

### 1.2. The varieties of hitchhiking

Not all is random during the gene shuffling of crossing over, as some genes move together in meiosis, rather like two sticky cards when shuffling a deck. A haplotype is a group of genes inherited together from one parent; the success of an allele of one gene in the haplotype carries along that individual’s versions of the alleles at other gene loci within the haplotype. This is gene hitchhiking.

I recently identified trait hitchhiking (Calvin 2017); it operates at the population level in an ecosystem. A trait’s alleles get a free ride around an amplifying feedback loop with no efficiency sculpturing; the trait becomes more common because its phenotype also had the habitat preference for being in the right place for an infrequent opportunity to have more offspring survive.

If the right place for a sudden expansion is in the brush bordering the grassland, then brush-favoring habitat preferences become more common. This also works for traits such as toolmaking which utilize the brush’s speckled shade for long periods each day; the alleles of those with toolmaking traits get a free ride around an amplification loop so that toolmaking ability also becomes more common in the core population. This is trait hitchhiking.

Because both types of hitchhiking can initially bypass the efficiency tests enforced by selective survival, they may be relevant to addressing some of the central problems in evolution that have proved intractable:

a. the doubling of our ancestors’ brain size in the first million years of the Pleistocene (Fig. 2);
b. promoting ultrasociality in *Homo,* another of the rare instances of eusociality; and
c. bursts of evolutionary extravagance such as our higher intellectual functions, for which selective survival explanations seem so inadequate.

*Figure 2: Brain size during the past 4 million years*

**Fig. 2.**
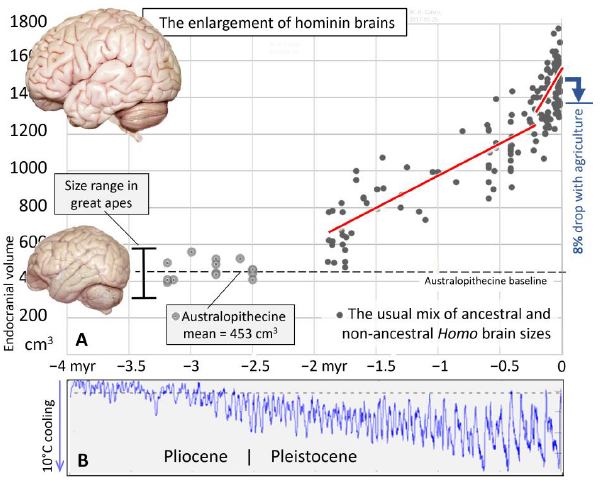
**A.** The evolution of preagricultural brain size as estimated from endocranial volume (excluding the more recent australopithecine-sized *H. naledi* and *H. floresiensis*). Australopithecines had brains little larger than those of the great apes (a bonobo brain is illustrated, to the same scale as the human brain). The average Australopithecine brain size (N=14) is shown by the horizontal dashed line at 453 cm^3^. After this long period of near-stasis, *Homo* brain size (N=161) increased 3.3 times, with the trend’s rate (red lines) appearing (but see Fig. 3) to triple in the last 0.2 myr, from 350 cm^3^/myr before to 1112 cm^3^/myr more recently. The blue down-arrow shows the 8% drop in endocranial volume with agriculture in the Holocene (Lieberman 1998). **B**. A proxy of ocean surface temperature (Lisiecki & Raymo, 2005) shows the increasingly deep ice age fluctuations in temperature, averaged worldwide over 57 sediment-core sites.

I addressed the evolution of eusociality earlier (Calvin 2017), together with the quantitative aspects (Fig. 1) of how amplifying feedback serves to repeatedly shift allele proportions (“gene frequency”). Here I begin the analysis of the steady brain enlargement of 460 cm^3^/myr since −2.3 myr (Fig. 3) and of the route to extravagance. There is an allele-amplifying feedback loop for a subset of meat-eaters that should generate extra neocortical space.

*Figure 3: Brain size in Homo sapiens and known ancestors*

**Fig. 3.**
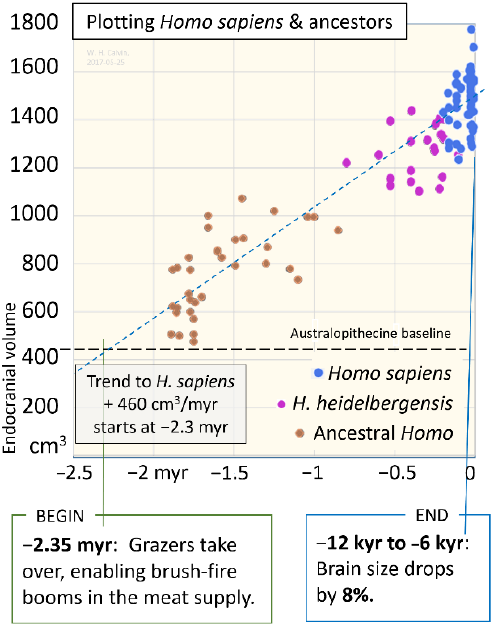
Separating *Homo sapiens* ancestors from nonancestors during the past 0.8 myr (from when *H. heidelbergensis* first appeared) clarifies the enlargement rate. The piecewise linearity suggested by Fig. 2A disappears if one excludes known non-ancestors. The dashed blue line intercepting the Australopithecine baseline at −2.3 myr is extrapolated from the least-squares fit to ancestral endocranial capacity (N=125) between −12 kyr and −1.9 myr, yielding a trend of 460 cm^3^/myr. The trend appears to begin at −2.3 myr.

## 2. The big brain problem

First I will analyze hominin brain enlargement, and then the ecological settings for hominin evolution, both in Africa and in the migration corridor north out of Africa for *Homo erectus*. Third, I will elaborate on the amplifying feedback loop and how it applies to hominin trait hitchhiking. Finally, I will address neocortical enlargement, reorganization, and “maps in search of a function.”

### 2.1. The social brain hypothesis

Though the inventions seen in the archaeological record demonstrate one aspect of intellect, the more common problems to be solved during hominin evolution were surely social ones during the evolution of food sharing and other cooperation. In the focus on how cooperation evolved, it was said that social group size should track brain size because increasing social complexity required a larger brain to monitor one’s cooperative exchanges with additional participants.

There are problems with interpreting this as causation. However well the social brain hypothesis may fit for primates in general, social complexity is no longer promising as a driver for the 3.3x hominin brain size enlargement. Maximum group size did not increase with brain size.

A band is usually several amalgamated family groups that travel together, the basic setup for food sharing beyond the family. It corresponds to nighttime group size, not to a foraging group, likely only part of a band. Once a band gets beyond a certain size, it tends to split permanently. Before agriculture, the maximum hominin band size (Marlowe 2005) remained stuck at a few dozen, about the same maximum that primatologists observe in gorilla, chimpanzee, and bonobo bands. If female exogamy requires exchanging adolescent females with a few other bands, that puts an upper limit on total individuals encountered at least once at perhaps 100-150. Yet everyday social life would only require tracking fair dealings with the dozens of one’s own band.

With settlements by −10,000 years, there were villages with a few hundred people with whom to cooperate (or not), all potential freeloaders to be monitored for fairness. Second-hand judgments of fairness, reputation spread via gossip, would increase in importance as group size expanded. Prosociality in the form of “social grooming” would eventually develop so that when a favor was asked for, there was already an established relationship to frame it, some history of give and take to recall.

By −7,500 years, group size was in the thousands; followed at −7,000 years by complex chiefdoms ten times as large (Turchin 2015). Towns, cities, nations, and other complex social networks, though abstracted from face-to-face social grooming (de Ruiter et al 2011), at least allowed travel without being attacked as an outsider, as would often be the case in the balkanized highlands of New Guinea.

Paradoxically, hominin brain size managed to enlarge 3.3 times with little if any change in group size. And then, when group size did finally increase, *Homo sapiens* brain size did not. It went down instead.

Even if the social brain hypothesis is true for humans—a big brain may well be a necessity if one is to keep track of a multitude of unique individuals—group size cannot be the *driver* of the 3.3x brain enlargement, which is the question addressed here.

The large-scale human prosociality, appearing within a few thousand years in a brain with only band-sized evolutionary experience, suggests that prosocial capability might have hitchhiked on some nonsocial evolutionary process before being tried out on the larger groups.

### 2.2. Hitchhiking in cortical enlargement

How do the early stages of a new neural function-say, the multiple mental countdown timers needed by short-order cooks-evolve via the selective survival of useful mutations, the best-known aspect of Darwin’s natural selection?

A bump-by-bump cortical enlargement, where each bump comes from a chance enlargement of an underlying functional map that is conserved by selective survival judging its usefulness, is what the standard evolutionary argument expects. Framing natural selection as simply selective survival sculpturing the neocortex piece by piece—a scenario that Finlay (2004) characterized as “Calvinist Cortex” though I would prefer to call it the Pointillist Presumption—has made it difficult to think about how more abstract levels of organization such as the higher intellectual functions could evolve to the point of a payoff.

It now appears that the developmental changes underlying brain evolution paint with a broader brush than the fine one portrayed in my pointillist caricature. Comparative mammalian studies (Finlay & Darlington, 1995; Finlay & Uchiyama 2015) show that the easy path to a local enlargement would be an overall neocortical increase, resulting in extra space for other functions that have no immediate payoff, e.g., if having more visual cortex were conserved by selective survival favoring those with better visual acuity, an instant side effect might be improved auditory discrimination. Or increasingly fine motor skills. Or room for tryouts.

Neocortical functions can thus hitchhike on an unrelated neocortical adaptation. There may be consequences to the broad brush, such as overloading the circulation or birth canal, but the variations may simply not be there to achieve a more efficient pointillist enlargement.

### 2.3. Hominin brain size during the last 4 myr

Scatter plots of hominin brain size (Fig. 2A) show four major features that any enlargement hypothesis needs to address, none of which are addressed by the social brain hypothesis:

1. a long period of near-stasis in the Pliocene in which brain size in the Australopithecines remained in the range for the extant great apes;
2. a fast 3.3x rise during the Pleistocene;
3. an (apparent) tripling of growth rate during the most recent 0.2 myr, and
4. a sudden 8% size reduction during the Holocene.

The late tripling of enlargement rate seen in Fig. 2A covers the same time span as *Homo sapiens*, which first appears in the fossil record at about –0.2 myr (McDougall et al, 2008). But the conventional scatter plot of Fig. 2A includes 36 data points from Neanderthals and Asian *Homo erectus*, now considered to be side branches, not human ancestors. To conflate what happens off the ancestral path while analyzing ancestral evolutionary sequence can confuse the issue, as appears to be the case with tripling and the earlier efforts using semi-log plots to fit exponentials to explain the slope increase at the high end.

In Fig. 3, these known non-ancestors are omitted, along with the earlier australopiths. Brain enlargement in *Homo sapiens* and its direct ancestors now appears regular at 460 cm^3^ gained every million years, using a least-squares fit from –12 kyr back to –1.9 myr, then extrapolated to cross the Australopithecine baseline at −2.3 myr.

There no longer appears to be a late spurt for *H. sapiens*; rather the question becomes one of why the rate remained arguably constant for so long.

### 2.4. Why did enlargement rate remain unchanged?

The lack of piecewise tendencies in the ancestral trend (Fig. 3) becomes a clue when the usual environmental drivers of adaptation are themselves so variable, so often. How could environmental challenges manage to consistently cancel out one another exactly enough to keep the brain enlarging at a constant pace? Evolutionary theory is seldom forced to address such a problem.

The lessons from complex systems analysis, however, would suggest that the brain’s observed enlargement rate may simply reflect just one of the many processes involved: the rate-limiting process is the slowest among all contributing to the enlargement process. The slowest supplier of parts limits the speed of an assembly line; if the slowest itself fluctuates, we will see rate variation in the output stream. But we do not see any change in output rate when the other suppliers fluctuate, so long as they do not slow so much as to become the new rate-limiting process.

While neural development is not a simple assembly line, there are surely more than a dozen developmental processes involved in cranial enlargement. If it is to continue to fit within the ensemble, only minor tweaks to a gene may be successful in avoiding developmental failure. Once the other gene loci have accommodated to the minor enlargement, it can be tweaked again (or, more likely, one of the ensemble’s other genes is tweaked). In this case, most of the time that germline evolution requires is not for natural selection to achieve a balanced polymorphism or go to fixation; it is simply waiting for another allele to be produced.

Once we realize that the enlargement rate is limited by the slowest of the processes involved, we are better prepared to identify it. The straight-line enlargement is what one expects of accumulation at a constant rate. What aspect of evolutionary change, from point mutation to haplotype formation in meiosis to demography to natural selection, is so reliably constant in rate, so insulated from severe environmental fluctuations?

### 2.5. Mutations in the germline: the role of the SNV

The typical successful mutation in the germline occurs when a cosmic ray collision in the upper atmosphere produces a neutron that, after its passage through the atmosphere, is moving slowly enough to knock out a single nucleotide without damaging so much nearby that the gene will not function. Gene repair processes insert a nucleotide into the gap but they make an occasional mistake, substituting a different nucleotide for the original, thus creating a single nucleotide variant (SNV, as in point mutation; SNP is a SNV present in more than 1% of the population) for offspring to inherit. The rate at which such repair errors occur may vary for certain genes, but overall the average mutation rate is usually assumed constant.

Some SNVs change which amino acid is added to the protein being assembled and may thereby create a new allele which in turn may cause the protein to fold somewhat differently, thereby tweaking functionality. A gene’s function can change rapidly when alternatives are already in circulation, as when the gene has alleles competing in a balanced polymorphism. When alternative alleles are eliminated in a selective sweep, the gene becomes “fixed” (four of every five genes lack alternative alleles).

Despite severe selection pressures, evolutionary change in a structure or trait may then stall until the next qualifying SNV. New SNVs occur at a rate of about once a year in human gametes (Kong et al, 2012), but the rate has not been constant during hominid evolution (Moorjani et al 2016), perhaps because the rate of errors in gene repair has altered.

Even though the interval between cosmic arrivals is random, their average rate should not drift. That is because of so many independent cosmic ray sources in the sky; nothing that happens on Earth will change their combined arrival rate in the upper atmosphere. This makes the constant cosmic ray background a candidate for the rate-limiting seen in hominin brain enlargement during the highly variable Pleistocene environment.

### 2.6. Achieving modern brain size

The line fit in Fig. 3 (V=1491 + 460T) intercepts the current year axis at 1491 cm^3^. The endocranial volume study of Lieberman (1998, Table 2) gives an average 1479 cm^3^ for pre-agricultural *Homo sapiens* and 1368 cm^3^ for Holocene skulls, a drop of 111 cm^3^.

This 8% reduction is unlikely to have caused retrogression of a new function—say, losing a new map or cancelling a recent reorganization of neocortical connectivity. That is because the concomitant gracilization in body shape beginning just before agriculture (Ruff 2006) suggests that maintaining selection for robustness (Brace 2000) was no longer keeping smaller individuals from contributing to the adult skulls available for our analysis; one consequence is that the average would drop.

### 2.7. Limitations on the fast track to the big brain

Two major candidates have been proposed for what could limit the rate of brain enlargement. First, the brain’s share of the blood supply nearly tripled, and this had to be at the expense of something else, likely prolonged digestion in a longer gut (Aiello & Wheeler, 1995). Second, increased head size (Rosenberg, 1992) had to be accommodated by widening the birth canal bottleneck and/or by slowing prenatal development and giving birth to ever more helpless infants.

There are physiological considerations as well, such as the “radiator theory” about effectively cooling a bigger brain during endurance running of prey (Falk 1990) that could slow down enlargement. However, even when slow enough to qualify as the rate-limiter, none seem likely to themselves produce a constant rate of enlargement in the face of major environmental shifts.

It would seem that producing new alleles is the ultimate rate limiter for brain enlargement; there may well have been situations where there could not be sudden spurts as new adaptations proved their worth if the feedback loop was already pinning enlargement rate to this limit. This suggests that gut reduction, birth canal adaptation, and brain cooling mechanisms were able to change more quickly than a new brain enlargement allele could break a fix—that they could adapt to the prior enlargement provocation during the period before another cosmic ray created an allele to further tweak brain size.

## 3. Ecological settings for hominins

We must consider whether brain enlargement is driven not by a function and its success but by a situation where brain enlargement is a continuing byproduct of a particular environmental situation. Functionality would change as a reaction to the enlargement.

With ecological situations in mind, consider the major events early in human evolution, starting with the bipedal woodland apes that added large herbivores to their diet (Pobiner 2016).

### 3.1. Foraging and confrontational meat scavenging

Australopithecines had prospered because woodland became much more common as African forests fragmented during the Pliocene cooling and drying (Cerling, et al 2013). The forest edges were woodlands; the more that the forest was parcellated, the more new edges there were, and the more prime habitat for the Australopithecines to exploit.

In a forest fringe newly experiencing direct, drying sunlight for some of the day, there were plant adaptations such as roots that store water, edible if pounded. The grass attracted large herbivores and their predators. Yet there were still enough trees for a new nest every night in the great ape tradition; trees were also a refuge when being chased.

Woodland was the Pliocene’s growth habitat and our ancestors were among the bipedal apes to seriously exploit it. They eventually were able to acquire substantial amounts of meat to supplement the edible-when-processed plants, likely via confrontational scavenging (Bunn 2001) at lion kills. The usual example of this type of interference scavenging (Shipman 1986) is the way a pack of hyenas can take away a kill from as many as three lions (Trinkel & Kastberger 2005) using harassment techniques.

I once suggested (Calvin 2002) that hominins could have used mobbing tactics to mimic this, but with the limited goal of quickly amputating a hindquarter and then departing; the lions would return to eat the belly fat rather than following the departing hindlimbs. Given how lions leave the limbs until last, contamination of their meat by intestinal bacteria would thus be minimal for as long as skin covered it.

Then a BBC (2011) video appeared showing three Dorobo hunters of southern Kenya, each carrying a long pole but not brandishing it, slowly walking up to a lion kill. One by one, the 15 lions fled, taking up positions behind nearby brush and watching.

The mobbing technique I had thought essential was not even needed. A lion can be cautious about novelties (if not an injured or underfed loner). There are now widespread reports of humans stealing from lion kills in Africa (Schoe et al 2009), some from wildlife management records a century old. They, presumably, did not learn lion behavior from watching staged spectacles.

### 3.2. Why sharp edges? Primitive amputation techniques

Scavenging has long been part of the hominin story. Shipman (1986, Shipman & Walker 1989) emphasizes the importance of disarticulating the carcass to be able to carry away meat. Here I examine a special case: the role of lion kills as a frequent source of sterile meat in large packages.

In comparison, the scavenging usually discussed has major limitations: infectious disease (for any meat that lions and scavengers have already chewed on, after feasting on abdominal contents) and infrequency (leftovers from the very top of the food chain). Once grazers evolved, their need for regular waterhole visits allowed lions to lay around in the brush near the waterhole, waiting to ambush them. The hominins could have done the same, awaiting an opportunity to steal hindlimbs.

Amputating a hind limb and leaving quietly could have been the big payoff for the sharp edge, not butchering more generally. Fast amputations would be consistent with hominin stone toolmaking being largely confined, for several million years, to producing sharp edges in quantity.

The Dorobo hunters may have used a large metal knife but there was nothing in their amputation technique that could not have been accomplished by two assistants forcefully stretching the skin while another used a sharp flake or fractured cobble to nick away at an exposed edge of skin to further encourage a tear. The pinch between thumb and forefinger needed to use sharp flakes may help explain the evolution of our precision grip.

Once inside, blunt dissection suffices to neatly separate muscles from one another, the result of which shows clearly in the BBC video after removal of the hindlimb—all of those smooth shiny muscle surfaces with no nicks from sharps. For attacking the joint capsule itself, sharp edges are again needed. For detaching the large muscles around the hip joint such as gluteus maximus, it would be helpful to use one of those long poles to lever the muscle taut, making it easier to sever near the tendons using only small sharp flakes.

None of these techniques would be important except for the considerable time pressure, what with hungry lions watching from behind nearly bushes. A supply of the little fingernail-sized sharps would need to be prepared in advance; it would not do to run out. Folding or rolling the sharp flakes up in leather, jeweler fashion, would have sufficed for transport. By −3.3 myr, hominins were producing sharp edges of some sophistication (Harmand et al 2015), 0.5 myr before *Homo* appeared.

The ubiquitous long poles seen around Africa today could have been another early invention; even without sharpening a tip by sanding it on a rock face, a pole can be used for clubbing birds and small mammals, and occasionally for warding off threatening carnivores. After that defensive experience, sharpening it for use as an offensive lance for thrusting must have been obvious.

### 3.3. Out into the grasslands

At −3.3 myr is when climate fluctuations first became noticeable (Fig. 2B), much earlier than −2.58 myr, the geologists’ most recent revision of the Pleistocene boundary. By −3.3 myr (Cerling et al 2013), the hominin diet had shifted from foods that originated in leaf-based C3 photosynthesis to emphasize grass-based C_4_, such as meat from grazers. At −3.3 myr, grazers were only 17% of the large herbivore taxa (Fig. 4, left window), so this change in diet comes well before the jump to 75% grazers at −2.35 myr. This is evidence that our ancestors at −3.3 myr were acquiring considerable quantities of meat by hanging around those essential waterholes.

*Figure 4: Proportions of grazer, browser, and mixed feeder taxa*

**Fig. 4.**
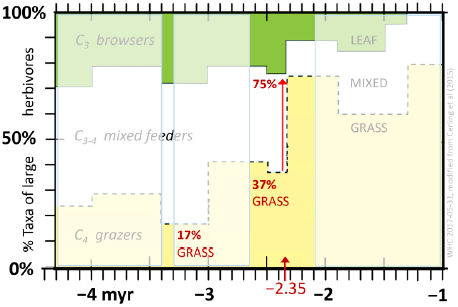
Relative proportions of browser, mixed feeder, and grazer taxa among the large herbivores in the Rift at the Turkana Basin, showing the major shift between −2.50 and −2.35 myr. Modified from Cerling et al (2015).

The waterhole strategy likely began with water close to woodland, then developed into overnight trips out to a treeless savanna waterhole. Volcanic eruptions near the African Rift tend to cover the land with a thick layer of dust that, with some rain, hardens sufficiently to preserve footprints and to prevent water from trickling deeper. Soil moisture then moves sideways in water sheets (the water table), often forming underground streams mere meters below the new surface. Elephants regularly in search of mud baths may hear the turbulence of the hidden streams and dig down toward the noise. The enlargement of the mud bath may allow a pool of water to form, attracting the large herbivores that specialize in grass and their predators. Eventually the grazers flatten the steep sides of the original waterhole, tending to plug up channels for water flow. Plant succession may not progress beyond brush before the underground stream is diverted elsewhere, giving such a waterhole a limited lifespan and few trees as a result.

Doing without trees to nest in at night and sleeping on the ground in the savanna demanded a level of social organization well beyond that needed for life in the forest fringe and woodland. Hominins eventually could sleep on the savanna when social evolution finally produced the sentry role: the duty to stay awake, sanctioned by the community if falling asleep or letting a fire go out.

### 3.4. The grass road out of Africa for Homo erectus

Early *Homo erectus* appeared on the East African scene by −1.9 myr, with a shoulder adapted for throwing projectiles (Roach et al 2013), suggesting that by that time they were making grazer kills themselves, likely only at a waterhole and not necessarily using techniques applicable to other large herbivores. With no further need of lions to serve as middlemen for acquiring grazer meat, they were able to follow a grass road out of equatorial Africa into Eurasia (Carotenuto et al 2016; Prat 2016).

The *H. erectus* shoulder at −1.9 myr suggests that spear throwing had been invented earlier, likely by adapting the ubiquitous long pole into a sharpened spear for thrusting, then into a javelin design balanced for throwing, with most of the weight in the front.

The initial targets were likely very big: the wall of grazers packed together at the waters’ edge; if it does not matter which one is hit, the situation makes minimal demands on accuracy. Developing the more difficult technique for hitting a single target animal is likely a separate evolutionary task (Calvin 1983, 2002)

The most direct route north, continuing this meat-dependent lifestyle of the East African Rift, would have been through the Afar Triangle, then up the Red Sea coastlines, north up the intersecting Dead Sea rift past the Sea of Galilee into Lebanon’s Beqaa Valley, then across Turkey’s rumpled landscape to Georgia’s Dmanisi at 42°N, between the Black Sea and the Caspian. They arrived at Dmanisi by –1.8 myr, their ancestors having covered 6,000 km, probably leaving behind relatives along the Mediterranean coast. The habitat near Dmanisi then was like the African savanna: grasses, arid conditions, and an abundance of large herbivores (Messager et al 2010).

Even when assuming this 6,000 km migration was packed into a 100 kyr time span (about 3,300 generations; see Fenner 2005), that is only 2 km per generation, a distance within their daily foraging range. That average advance in 30 years was likely achieved by a seasonal back and forth as the easterlies’ rain band moved north and south; their campsites would have gradually moved north because the herds there were more naïve and therefore easier to hunt.

### 3.5. Sustained migration and the confining corridor

What took *Homo erectus* from the trade-wind tropics to a continental climate with harsh winters? Certainly not planning. Nor the realization that their drought was part of a longer-term trend in precipitation and temperature that warranted getting out of Africa. There is no evidence that the first human dispersal about −1.8 myr coincided with any major environmental change (van der Made 2011).

There is a more elementary migration process that works well—but only for hominins who had already adapted to Rift ecosystems and then eliminated the middleman for meat. When a herd of large herbivores had never encountered a two-legged hunter before, they were likely easy prey. A hunter might slowly approach a naïve herd if remaining upright like a tree and never crouching down like a lion. After being hunted a while, the remainder of the herd would become so wary that hunters would move on to the next naïve herd, centered on another source of drinking water.

In a north-south corridor often bordered by deserts or bodies of salt water, the hominins cannot disperse east or west, and south is the direction of depleted and wary herds, taking a few years to recover its numbers and for memories of two-legged predators to die off. And so, like a burning fuse or a propagating nerve impulse, they advance to the next hunter-naïve herd farther north on the grass road.

Some might have spread north to populate the Mediterranean coast. By going east along the northern Sinai at 31°N, they would encounter the Dead Sea Rift and resume the northbound burning-fuse migration with the grazing herds there. The strengthening westerlies meant that the migrating bands would encounter even more east-west dispersion opportunities in the Fertile Crescent. Eventually winter weather would be encountered and the hunters would have needed some non-African adaptations to stay with their meat source for the winter rather than seasonally retreating. The nomadic Inuit and the nomadic Sami are a modern example of such a meat-dependent lifestyle.

While African flora and fauna make some inroads into the Levant, the edible plant possibilities farther north would be novel, requiring generations to adapt food preparation techniques for removing plant toxins successfully to the unfamiliar plants en route. Living this way is eventually possible but it would not support very many, compared to meat. While the diets of stay-behinds would be a mix of strange flora and fauna, the diets of those in the corridor would be familiar: sterile, toxin-free meat.

While meat may have fueled the human trip around the world, note that the travelers’ diet is not the average diet: what they would have eaten (see Schoeninger 2012) if there were more choices than just the next naïve herd. Nor are such large herbivore techniques a model for the development of carnivory in hominins; in Africa, the evidence is that they started small (Ferraro et al 2013), perhaps using long poles to club animals hiding in the grass as even some African children do today.

## 4. Amplifying feedback speeds evolution

As summarized in Fig. 1, boom-and-bust dynamics creates a “gene pump,” a way of repeatedly combining selective reproductive opportunity (founder effect) with a subsequent squeeze that can quickly produce a substantial shift in main population’s allele proportions (“gene frequencies”), what would otherwise take many millennia of selective survival operating on the fringe fraction of the population. The characteristics of such amplifying feedback fit well with the major characteristics seen in Fig. 2A and 3.

### 4.1. Why did enlargement begin at –2.3 myr?

It is apparent from Fig. 2 that brain size doubled in the first million years of the early Pleistocene, an extraordinary event in evolution. The fitted line for ancestral *Homo* in Fig. 3 extrapolates back into the 0.5-myr-wide fossil crania gap (though they go back to −2.8 myr, *Homo* specimens earlier than −1.9 myr are scarce: two mandibles, one maxilla, one frontal bone and few teeth).

This ancestral trendline crosses the Australopithecine baseline at −2.3 myr. Why might enlargement start just then, after millions of years of little growth while the Pliocene climate cooled?

The number of large-grazer taxa at Turkana in the Rift Valley doubled between –2.50 myr and – 2.35 myr (Cerling et al, 2015). In that same 150 kyr period (Fig. 4), mixed-feeder taxa dropped by two-thirds and browser taxa dropped by half. Specialization for efficiently eating the plentiful grass all day was a response to climate change. Yet there was still enough dense brush in places to allow an amplifying feedback loop to operate after a lightning strike back in the brush. This exposes some hidden aspects of evolution.

### 4.2. Brush fires provide a boom time for grazers

There is a high rate of lightning strikes atop the mile-high savannas of East Africa and South Africa. Both grass- and brush fires result and a large area can burn in the dry season. New grass is soon pushed up from below. A grass-only fire would have little effect on herbivore populations, as would a grass fire that made inroads into flanking brush. But if a burn scar back in the brush has a path connecting to the core population, large herbivores may discover the burn scar’s uncropped young green grass.

Provided that the burn scar contains enough drinking water, this allows brush-fire booms (Fig. 5) to commence in the grazers, and only the grazers: the burn scar diminishes resources for browsers; for mixed feeders, the temporary grassland had already been part of the territory determining the carrying capacity (they would merely be switching to their less nutritious option, not booming).

*Figures 5 & 6: Boom and bust migrations power the feedback loop*

**Fig. 5.**
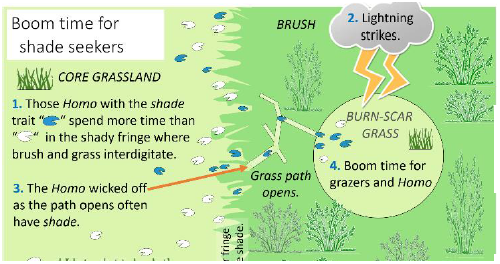
The boom-time setup for feedback.

But for the specialized grazers, which became 75% of the large herbivore taxa by −2.35 myr, intact brushlands were not a resource and grass availability controlled the carrying capacity, together with the predators. With an auxiliary grassland to discover back in the brush, one expects a grazer population expansion over a few decades as so many of the excess-to-replacement offspring manage to grow up there and reproduce themselves, what with more resources than they can consume. This boom time might have continued for as many as ten grazer generations. The feedback might have played a role in the fast conversion of mixed feeders to grazers at −2.5 myr.

If this auxiliary grassland wicked off fringe-favoring hominins who also knew how to acquire meat in large packages, subpopulation booms similarly became possible in the *Homo* lineage, though there is only enough time for grandchildren before brush returns to slowly lower the grazer carrying capacity in the temporary grassland.

Earlier I quantitatively analyzed (Calvin 2017) the brush-fire cycle behind the grassland’s brushy frontier to see how much the feedback loop shifted allele proportions (“gene frequency”). A selective expansion (Fig. 5) coupled with return migration (Fig. 6) can produce a substantial gene shift in the core within relatively few generations. And feedback can operate without concomitant selective survival.

**Fig. 6.**
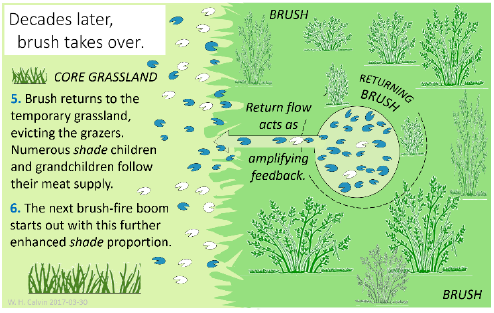
The bust produces allele-amplifying feedback to the core, an example of non-random gene flow.

It might be asked why the brain enlargement is linear if the feedback loop can create exponential rises. The exponential rise is in the allele ratio (Fig. 1); when fixation is achieved, it stops. Were the step size constant rather than escalating, it would make little difference, given how quickly fixation can be achieved with a feedback loop, well before the next arrival of a cosmic ray can create another allele. The linear rise in brain size is attributed here to everything else being faster than the average time between cosmic rays, and that that rate-limiter did not change its rate over 2.3 myr.

### 4.3. What makes the Rift Valley ideal for feedback loops?

The East African Rift and its northern extensions are all spreading centers. Beneath them, the hot lithosphere is closer to the surface; the heat expands the crust, it domes up, and then parallel cracks may form, allowing more independent movement in the future. The crust between two parallel faults can drop down as a wedge-shaped block (Fig. 7) when lateral support is removed by spreading.

*Figure 7: Dropping down into a rift depression*

**Fig. 7.**
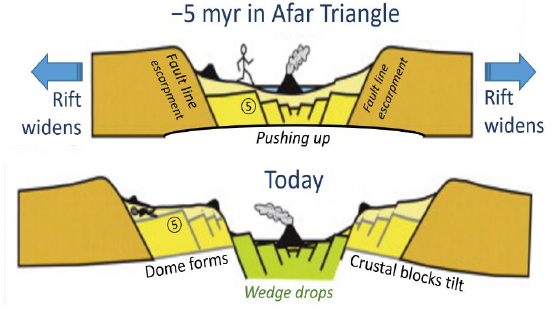
The Rift valley cross-section at the hominin fossil sites in the Middle Awash of Ethiopia. The diagram shows the change over 5 myr; the crumpled stick figure shows where ancient bones might be found today. Modified from Fig. 16 of Bailey et al (2010).

This produces a characteristic appearance (Fig. 8 & 9): a long valley, steep escarpments and, behind their ridgeline, one may see flat country at much the same elevation. Erosion may modify the valley, but this is not the usual valley carved by the flow of water, partly filled in by layers of sediment. The depression may be filled by fresh water, as at Lake Turkana, Lake Tanganyika, and Lake Malawi. If the long wedge drops far enough, salt water may find its way into the valley, as at Lake Assal in the Afar Triangle. A dramatic example of a flooded rift is the long, narrow Red Sea with its parallel escarpments behind the coastal plains. This intersects the Dead Sea fault leading up the Jordan River to the equally depressed Lake Tiberas (aka Sea of Galilee; the Dead Sea bottom is at −735 m; Tiberas’ is at −739 m relative to modern sea level).

*Figures 8 & 9: The Rift Valley setup for a feedback loop*

**Fig. 8.**
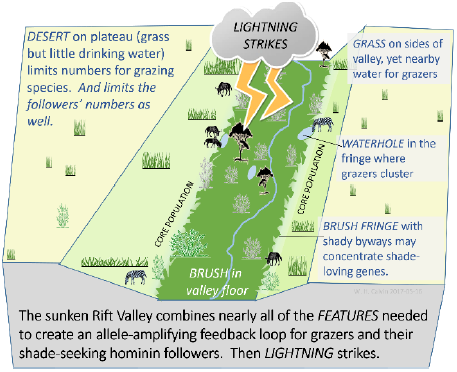
The Rift Corridor features that affect grazers and their followers.

**Fig. 9.**
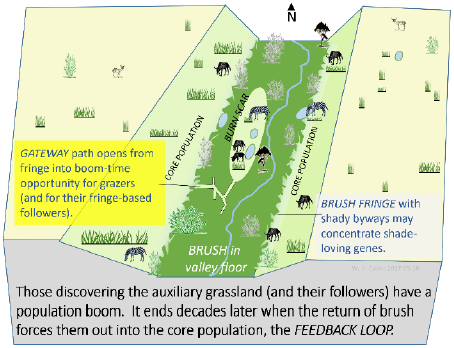
Corridor-based boom-and-bust feedback; the return path can shift gene frequencies even more readily than when the Rift is wider.

To understand why grazers cannot utilize the grass atop the escarpment and so enlarge their population and that of migrating *Homo*, consider what happens to desert rainfall near spreading centers. Desert surfaces often do not retain rainfall; soil moisture tends to descend until reaching an impenetrable ash layer. This makes streams and waterholes rare on the plateau.

This sloping water table (Fig. 10) may come out of the escarpment as a waterfall or lower down as a spring; examples are Ein Gedi and Jericho in the rift of the Dead Sea. The valley floor thus collects water funneled in from adjacent desert; it may support brush or even trees, as near the Jordan River. But slightly uphill, there is often a brushy fringe interdigitating with grass.

*Figure 10: Groundwater emerges from the Rift’s walls*

**Fig. 10.**
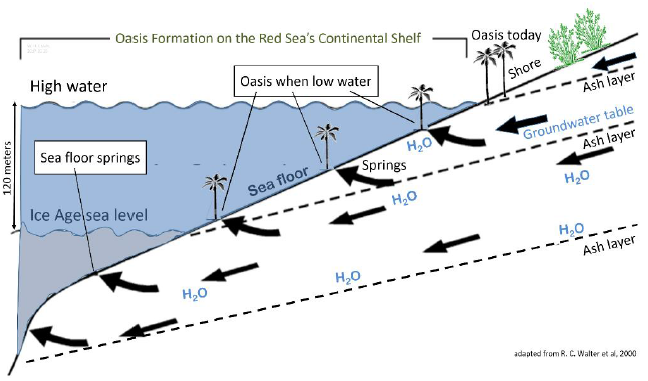
Ashfalls from local volcanos create a series of impermeable layers along which rainwater flows downhill, with hydrostatic pressure developing. Sea level is currently 130 m higher than in glacial extremes, drowning many of the springs around Africa’s margins. At low sea levels, Red Sea coastal plains would have had a series of oases with grass and brush for large herbivores. They would have served as refugia and migration routes for hominins during the cool-dry periods in Africa. Adapted from Walter et al (2000).

Higher, and it may be grass with occasional isolated bushes but no drinking water for grazers. Grass and the occasional bush may continue over the top and back a considerable distance. Mixed feeders such as elephant and eland can utilize it if they can get enough water from the leaves, but the grass specialists stay within commuting distance of a reliable water source. Grazers come down to water in tightly packed herds every other day in a manner that mixed feeders need not, making grazers easier to hunt than other large herbivores.

### 4.4. Migrating in confining corridors

The requirements of big grazers may be the key to understanding early *Homo* migrations. Indeed, the earliest *Homo,* at −2.8 myr (Villmoare et al, 2015), was recently found in the Afar triangle at 11.3°N, a convenient location for getting caught in a burning-fuse migration up the Red Sea coastal corridor 2,100 km to the Mediterranean at 31°N, about the distance between the Canadian and Mexican land borders. Most such migrations would dead-end on a gap without freshwater resources for the grazers. With only enough freshwater for the hominids, the gap might still be bridged using offshore resources.

Most importantly for feedback considerations, a long green trench through a desert limits the population of the boom-prone grazers, which in turn limits the numbers of their followers. As noted, hominins following grazing herds north in the Rift should result in a burning-fuse Out of Africa migration. The Red Sea coastal plain could have become a major migration path (Walter et al 2000; Beyin 2013), bypassing the Sahara’s arid barrier.

Once a narrow corridor is encountered, the dynamics of evolution can change. Such a northerly migration corridor is capable of hosting amplifying feedback, operating repeatedly on a migratory band over many generations for 5,000 km of confining corridor.

The Red Sea is not noted today for a connected string of grazer-accessible waterholes extending up its coastlines, but sea level is currently high. In glacial times, it was as much as 130 m lower, uncovering a series of springs that had previously been pushing out fresh water below sea level despite the hydrostatic pressure (Faure, Walter, & Grant 2002); some had such abundant flow in recent centuries that the sea surface above the submerged outlet had so little salt taste that sailing ships would restock their drinking water by dipping barrels into the upwelling.

So, along many of Africa’s continental margins, there would be brush, fresh water, and grass for grazers and their followers at a time when much of Africa’s interior was exceptionally inhospitable during the cooler (and thus drier) glacial-era climate (Faure, Walter, & Grant 2002). Coastal plains should have repeatedly acted as refugia for our ancestors, once they could acquire meat without lions to act as a middleman, an early case of disintermediation.

Figures 8 and 9 summarized the situation in the Rift Valley more generally, showing a desert plateau above the escarpment, with its lack of water. Substituting a salt water barrier for the eastern escarpment reduces both prey and predator populations by half (Fig. 11). But with the burn scar population not changing, this can double the effect of a single brush fire on core gene frequencies.

*Figure 11: The confining half-corridor*

**Fig. 11.**
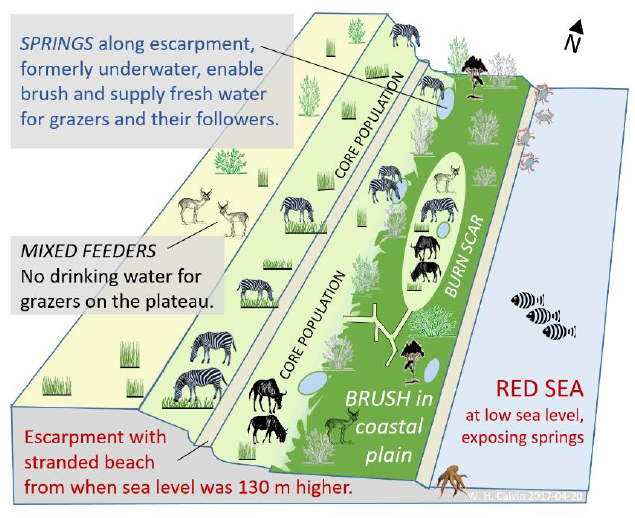
The confining (for grazers and followers) corridor. The Red Sea rift’s coastal plains at low sea level provide an even better setup for feedback enlargement than the two-sided Rift Valley of East Africa.

### 4.5. Migrating bands in a confining coastal corridor

Offshore coral reefs within wading distance made seafood a dietary alternative along Africa’s coast; freshwater catfish and turtles were already utilized by −1.95 myr at Turkana (Archer et al 2014). Indeed, some evolutionary aspects of our physiology such as the diving reflex are difficult to explain without periods when our ancestors exploited offshore resources (Morgan 1982).

Springs dotted along the coastal plain at frequent intervals could have made a coastal refugium into a migration path. Were there a gap without enough water for grazers, northerly progress would have halted, were meat from grazers the only hominin tactic. However, if they began using the ample seafood offshore, the only remaining barrier would be enough water for the hominin band itself, perhaps from small waterfalls in the escarpment such as Ein Gedi. Again, over-harvesting of local seafood would force them to search northward where they might eventually find another grazer herd.

Such a confining corridor migration poses a problem: contact with other bands is needed to continue the practice of female exogamy. In the broad parts of the Rift, it is easy to imagine a band’s annual circuit providing a half-dozen opportunities to meet up with nearby bands, in places where resource density allows seasonal opportunities to feed larger groups for a short time. But in a one-dimensional coastal corridor, bands are spaced out by the aftermath of a band’s passing: wary herds and depleted offshore resources. Bands of a few dozen move on when resources become insufficient for the full population of the band. However, the corridor might remain adequate for small groups, both southbound and northbound, thus achieving female exogamy.

If mixed feeders also occupy the grassland (Fig. 7), they squeeze the size of the grazer population. Because mixed feeders can move on to leaves on the plateau when grass is depleted, they have more options than the efficient grazers.

### 4.6. Failure modes and non-ancestral Homo

Keeping the grazer population relatively small is how the returning gene flow from the defunct auxiliary grassland (Fig. 6) can make a larger shift in the total population’s gene frequencies (Fig. 1).

Let the hominin population become large, as did agriculture’s boom, and the impact of a single meat-boom loop becomes insignificant in a manner familiar to Malthus: after all, the carrying capacity of an auxiliary grassland (the average meat boom) does not scale up as the core population increases. Thus effect size falls.

As with chemistry’s autocatalytic processes, this allele amplification loop has failure modes. Had grazers been hunted to extinction, browsers and mixed-feeders would still have provided a meat supply for *Homo*, new alleles would have continued to form, shade might concentrate certain traits, lightning strikes would continue to create grassy burn scars for them to discover, but there would have been no meat boom to run the allele amplification loop.

Partial failure occurs if migrating into a region with few sites suitable for forming a brush-fire feedback loop (deep brush adjacent to grassland). The non-ancestors omitted from Fig. 3 are plotted separately in Fig. 12 together with the ancestral *H. erectus* (older than the −0.8 myr appearance of our ancestor *H. heidelbergensis*). It is apparent that since −0.8 myr, the non-ancestral Asian *H. erectus* created the sag in the middle of the combined data of Fig. 2A, creating the appearance of tripling enlargement rate by lowering the earlier slope.

*Figure 12: Lags in enlargement in nonancestral Homo*

**Fig. 12.**
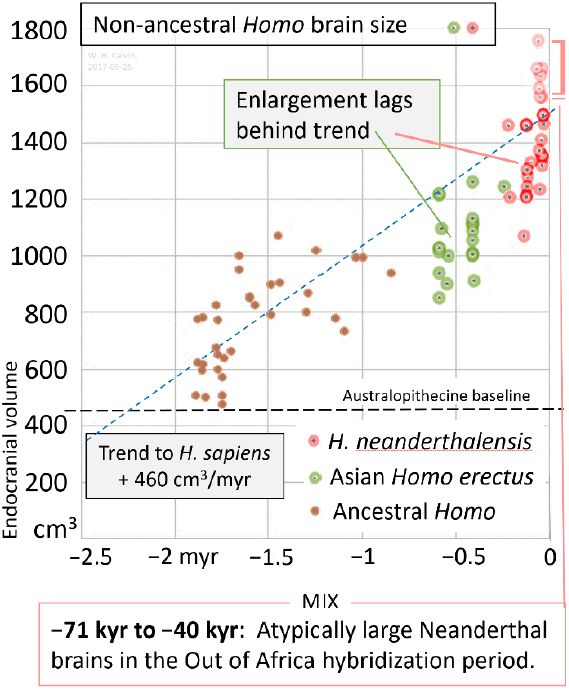
The non-ancestors removed from Fig. 3 are plotted, showing lags in enlargement for Asian *Homo erectus* and Neanderthals.Dashed lines as in Fig. 3. Plotting the omitted brains shows that both non-ancestors (N=36) lag the ancestral trend line, taking as much as 0.5 myr longer to achieve a given size. Note that the Neandertals have a substantial brain enlargement between −71 kyr and their disappearance at −40 kyr, just after the major African migration into Europe at −45 kyr. The seven largest Neanderthal brains (bracketed, >1550 cm^3^) occur in that period, matching the top end of the *Homo sapiens* size range in Fig. 3.

The horizontal separation of the green Asian *H. erectus* data points from the ancestral dashed blue line in Fig. 12 shows that they took about 0.5 myr longer than our ancestors to attain the same brain size. Too much grassland, as in the steppes of Central Asia, dilutes the loop increment because there is, relatively, little brush border. Similarly, migration into forested Europe would have made suitable sites (deep brush bordering grasslands) infrequent. However, brain size would continue to increase, just more slowly, if there were a continuing stream of new bigger-brain immigrants from Africa.

A briefer lag in enlargement is seen in the Neanderthals; if one ignores the seven Neanderthal brains above 1550 cm^3^, most red data points are well below the dashed trendline for *H. sapiens* ancestors imported from Fig. 3. The seven are all from the period between −71 kyr and the final, widespread cultural disappearance of the Neanderthal’s Châtelperronian and Mousterian artifacts (Higham et al, 2014) at −40 kyr.

The Out of Africa at −70 myr for *H. sapiens* is usually characterized as spreading up the Levantine Corridor, turning east in the Fertile Crescent to spread along the Silk Road route, with a branch along the west coast of the Indian subcontinent. The biggest Neanderthal brain, 1745 cm^3^ at −63 myr, is from the Amud cave just inland of the Mediterranean coast in northern Israel.

This late Neanderthal growth spurt, so much faster than the trend to *H. sapiens* over 2.3 myr, suggests a different enlargement regime in the Neanderthal’s final period, such as acquiring bigger brain alleles during mating encounters with *H. sapiens,* much as we (with the exception of Africans) acquired our collection of Neanderthal genes (Pääbo 2014). This shows how rapidly brain enlargement can occur if there is not the wait for another allele to be produced; indeed it suggests that some of the “enlargement ensemble” of genes move together as a haplotype.

### 4.7. Trait hitchhiking in the feedback loop

For both prey and predators, the boom-and-bust feedback loop can repeatedly shift the core’s gene frequencies toward those of the brush explorers. Hitchhiking permits deleterious traits to bypass editing by selective survival, perhaps creating an increased prevalence of myopia in the population. Any brush-relevant allele could benefit from this amplifying feedback (Calvin 2017), so long as its phenotypes concentrate near where opportunities can open up back in the brush. For example, myopic individuals who made poor foragers would have tended to take on close work in the shade, such as pounding, chopping, and creating sharp edges for those better at hunting and gathering. Myopia alleles would become more common in the core population, after many rounds of a brush-fire feedback loop.

If behavioral versatility concentrates in the fringe, this amplifying feedback loop would have affected brain enlargement. Given equal body size, the versatile omnivores have larger brains than those in taxa with one specialized food strategy. For the feedback loop setup, those versatile enough to skillfully hunt both grass-eating grazers and leaf-eating browsers would spend more time in the brush, making them more likely to be there when a gateway opened to the hidden grassland. Those specializing in open-terrain confrontational scavenging (Bunn 2001), or running an herbivore to exhaustion (Lieberman et al 2009), might miss the opportunity.

However, other types of versatility would be more commonly affected.

### 4.8. Clustering in the shade

The immediate border of the grass road may be a shoreline or a hyper-arid zone but the usual border is brush, providing shade when seated. The border usually has a ragged edge, with narrowing paths of grass that terminate in a brushy dead end.

Who needs to hang out in the brush’s speckled shade? Most commonly (Calvin 2017), mothers holding an infant on a hot day, at least if she wants to work with both hands, as when pounding or chopping. Small bodies have a higher surface-to-volume ratio than a large body, and so can change core temperature much more quickly. Just as a hairless infant requires heat from an adult’s body when it is cool, so avoiding heat stroke means that the infant needs skin-to-skin contact with a large heat sink, one with more evaporative cooling capacity. If the infant cannot be handed off to a large-bodied helper (a child may not suffice), then in order to park the infant, relocating to the shade may be required to avoid heat stroke.

Secondarily, there are activities that require sustained attention where shade is sought in the brush fringe by those spending hours attempting to start a fire, those creating sharp edges, and those chopping or pounding roots. While almost everyone has basic foraging skills, individuals who are also able to do such focused tasks are more versatile, which has interesting consequences.

### 4.9. Assortative mating

The clustering of these more versatile adults, while others are away hunting and gathering, provides a setup for assortative mating. Some children of such matings will be able to sustain attention even longer, increasing their versatility and eventually their brain size.

A modern example is the way in which college admission tests serve to cluster those with a long attention span, and at an age when mate-finding is peaking. As with the heritability of stature, only a minority of the resulting children will match or exceed their parents because of regression to the mean. But some will.

As it is usually conceived, assortative mating merely increases the spread, not necessarily the mean. Offsetting the extra tall offspring are the extra short offspring produced by the below average parents, more so because fewer of the tall would be mating randomly with the short. However, this might be changed into a trend toward even taller if there is a filter—say, too short to thrive—that keeps the new low end from reproducing via juvenile mortality.

College admissions assortative mating changes in the genome are not considered significant on the time scale of 35 years (Conley, et al, 2016) but evolution usually operates on a far longer time scale than just one generation. The observed brain size enlargement, 460 cm^3^/myr, only changes brain size by about 0.16 cm^3^ in that period, a mere 0.016% per generation for a 1000 cm^3^ brain.

Brain size is as strongly heritable as stature (Witelson et al, 2006), so bootstrapping brains to a new normal may be done by assortative mating; adding a feedback loop may make it happen much more quickly. Note that an evolutionary arms race is not needed to continue the enlargement and there is also no need to keep proving the worth a slightly bigger brain via its contribution to survival rate. Trait hitchhiking per se may bypass selective survival (though countering processes may still be affected).

## 5. Building up versatility

There is more to enhanced functionality than just the overall enlargement of the brain. Reorganization en route to the human brain also played an important role (Teffer & Semendeferi 2012), both for relocating maps and for interconnectivity patterns.

There is now an important insight from inferred blood flow that echoes Falk’s “radiator theory” (1990): as hominin brain size increased, the internal carotid blood flow (inferred from foramen diameter) increased almost twice as much as that expected for oxygenating the extra number of neurons (Seymour et al, 2016). This suggests that the brain began performing more neural computations simultaneously.

### 5.1. Free rides and the many ways to hitchhike

Three hitchhiking examples have been mentioned here:

1. Some neocortical functions should improve with natural selection for performance in other unrelated neocortical functions.
2. Hauling along the whole haplotype.
3. Trait hitchhiking at the level of demography.

The first example of a free ride was Darwin’s (1859) analysis of conversion of function. His example was the fish’s swim bladder, used for regulating buoyancy via expanding an air sack with blood gases. It was then converted into an organ specializing in gas exchange between blood and inhaled air. Soon thereafter (Dohrn 1875), succession of function became an example of how a novel function can be promoted in evolution: just repurpose an earlier structure.

Or duplicate it and then repurpose one of them, which is what the many visual hemifield maps (Wang et al, 2015) in neocortex suggest. A simple developmental soft-wiring rule (“Neurons that fire together wire together”) may suffice to create the retinotopic visual field maps in neocortex, given an infant’s overly broad connectivity and enough visual experience in the right time frame before the less utilized connections are pruned.

### 5.2. Maps in search of a function

An example of the parallel processing that might be keeping the brain so busy: in each hemisphere, the human brain now has at least 25 retinotopic maps of the visual field (Wang et al, 2015; Mackey, Winawer, & Curtis 2017), not just the ones in textbooks. A moving object is thus analyzed simultaneously by many specialized processes.

Early in life, a neuron is often observed to have many more axon terminations than the counts from a juvenile or adult (Lund et al 1977). The pruning was initially seen as a necessary step for myelination, making space in the corpus callosum for the myelin wrapping. But pruning also allows the developmental soft-wiring rules to sculpture maps and other associations, even allowing different maps to overlap. That creates an opportunity to randomly self-organize overlapping representations and try them out, perhaps yielding a novel cognitive capability defying description. Synesthesia is likely an example of such “Maps in search of a function.”

Note that the individual variability in how maps overlap, and occasionally give rise to new neocortical specializations, provides a new level at which natural selection can operate.

Humans have 180 neocortical areas in each hemisphere whose boundaries have sharp changes in cortical architecture, function, connectivity, or topography. That is somewhat more than in a monkey (Glasser et al 2016). A candidate for a unique add-on: Orban & Fausto (2014) describe a tool-use specialization that is located in the human left anterior supramarginal gyrus, near the primary sensory representation for the hand. This area, which cannot be found in the monkey, seems devoted to the execution and observation of manual tool actions.

### 5.3. Ballistic movements: when feedback is too slow

In addition to behavioral versatility building up during hominin evolution, there were major improvements in the hand-eye coordination for the “get set” ballistic movements, such as the increasingly delicate hammering needed for accurate throwing from ever greater distances and for the fine serrated cutting edges of Achulean-style tools (Dibble et al, 2016) and the sharp flakes that are produced from such a core.

Ballistic actions may require a complete plan before fast movement begins (saccades, clubbing, hammering, throwing, spitting, kicking). That is because any feedback arrives too late to change the fastest part of the action: the 120 msec reaction time for correcting forelimb perturbations in humans is about the total time for a dart throw, and so any reflex correction arrives after launch. Flinging, however, is often slow enough not to need the complete plan in advance.

The need for more and more precise timing of forelimb movements in throwing and hammering could, it was suggested using Law of Large Numbers reasoning (Calvin 1983, 2002), be accomplished by a wider and wider synchronization of neocortical neurons immediately before launch.

### 5.4. Syntax: a developmental role for gossip in the shade

There is bootstrapping across generations as the larger-brain, more versatile adults provide structured sentences for young children to hear during the sensitive periods for tuning up intracortical connectivity’s “soft wiring” of early childhood.

Gossip (see Dunbar 1996, 2004) would be the most frequent source of structured sentences for young children to hear. Structure is not needed for the short sentences of protolanguage (what the modern two-year-old is mostly speaking) but it becomes essential for disambiguating longer sentences in their third year and for them to subsequently produce long sentences in a form whose meaning may be readily guessed by listeners.

Yet what was the evolutionary foundation that encouraged verbal gossip, with all of the attention paid to “Who did what to whom, where, when, and with what means?” It would eventually be useful for reputation, once group size exceeded the number that one can judge for oneself, but a more immediate driver would have come from female exogamy. Where female mate choice is possible, the reputation of a male becomes important for a newly-arrived adolescent female who is uninformed about all of her potential mates.

Facial expression, arm gesture, gaze, and pantomime can all be used to gossip; indeed, in modern sign language, long sentences are facilitated by pointing to one corner of an imaginary tray to define a noun phrase (“The tall blond man with one black shoe”), later pointing at that corner when verbal gossip would instead use a personal pronoun to point back. Once invented, ehe imaginary tray provides four repositories, easily reusable.

Yet this is laborious when adult hands are busy with shade tasks or infants to hold; facial gesture and direction of gaze would have to carry the communication load. Gestural communication also takes eyes off the task and is usually ineffective after the quick tropical sunset. Great apes rarely monitor one’s direction of gaze at objects, nor do they respond to pointing by orienting or moving; australopithecines might have shared those tendencies. Gesture seems an awkward way to start, however useful it may be in supplementing verbal strings or communicating during silence.

The verbal version of gossip is, however, confined to a linear string of arbitrary sounds and so it needs other ways to point back to where a noun phrase was created. For long sentences, each dialect has local structuring conventions (grammar and syntax), enabling a listener to quickly disambiguate the longer complex strings such as “I think I saw him leave to go hunting.” The listener must sort out each of the four nested verbs in that sentence, determining which noun goes with which verb.

### 5.5. The inverted pyramid for language development

Long-sentence structuring is the fourth in a series of self-organized tune-up stages that a child normally experiences in its first three years, given enough overheard utterances to absorb. Infants initially self-organize for the briefest of the speech sounds they routinely encounter (Kuhl, 2004). Later in the first year, the phoneme-competent infant then categorizes the concatenations of phonemes and learns many new words every day. They are also busy forming concepts (Bloom 2001); in the second year, categorization efforts appear to span strings of a few words.

In the third year, categorization over a still longer time span allows the child to first understand and then generate long sentences. A fifth tune-up period in the preschool years comes as they notice patterns in a group of many sentences, such as a scenario. The child may then demand bedtime stories that match Aristotle’s template: a plot with a proper beginning, middle, and end that is nonetheless short enough for the recipient’s attention span to encompass.

So five levels of organization have been built, just in the preschool years. Later, additional levels are built for analogy and parable; some can be utilized for the comparisons and ordering needed for complex thought and rationality (Calvin 2004).

If this *H. sapiens* pyramiding sequence for language development existed in some form in earlier hominins, any verbal structured version of gossip would be routinely overheard in the shade by nearby infants and supervised children, providing examples for their tune-up processes. Then comes an extraordinary bootstrap: the child could then grow up to be even more proficient than its exemplars, those gossiping adults who had learned language somewhat later in life. Experiencing complexity earlier in the sensitive period for category formation likely improves adult performance; certainly, failure to provide an infant with early exposure to complex examples, neither spoken nor sign, may result in a modern child growing up with severe limitations in intellect.

One can imagine a group losing their more language-productive adults so that a subsequent infant would not hear enough complex examples to categorize long sentences. The band’s ensuing period of reversion to protolanguage (and likely diminution of complex thought) might end when a newly arrived adolescent female from another group provided structured examples for local infants once more. Culture not yet backstopped by genetic-based instinct is prone to loss, requiring reinvention or travelers to reinitiate.

While there is little in this protolanguage-to-language example (Calvin & Bickerton 2000) that is exposed to selective survival, the shady setting for child care while working bimanually provides access to selective expansion’s boom-and-bust loop.

## 6. Summary and Conclusions

In paleoanthropology, when estimating how important an evolutionary driver is, we must take account of fast tracks, distinguishing between the more general influences (say, when travel is merely regional circulation for foraging) and those relevant to situations where *relative* speed becomes important.

Two types of fast track have been described here that change allele proportions. The coastal corridor not only serves to channel but it limits population size and promotes first-cousin matings, speeding evolution. Second, the repeated operation of the brush-fire feedback loop serves to accumulate small tweaks much more quickly than otherwise.

The feedback loop itself has a need for specialized grazers in order to produce a meat boom after a brush fire; a need for some isolation spanning generations; and a return path to the core when brush takes over again. For trait hitchhiking around the feedback loop, it needs heritable traits that concentrate somewhat in the brush’s fringe with grassland, such as shade-seeking for fire-starting, toolmaking, food preparation, and the communal care of infants. This clustering permits assortative mating of, for example, the more versatile to increase mean brain size.

Brush fires happen so frequently, and the time it takes for a mutation to go to fixation is so short given amplifying feedback, that enlargement appears to be rate-limited by how often cosmic ray neutrons strike a relevant gene in a way that results in a new allele able to compete with the fixed gene.

Here I have used versatility as a serious possibility for the enlargement driver but the amplifying feedback could instead operate on other heritable traits affecting brain size that concentrate in the brushy fringe—suppose, for example, that brain enlargement was linked instead to myopia or to strong handedness for ballistic movements.

The present analysis establishes a rationale for all six major details of hominin brain enlargement.

1. *Why Australopithecine brains did not enlarge*: even if they were regularly acquiring meat and populating burn scars, there were not enough boom-prone grazers and so no booms in the meat supply.
2. *When enlargement began:* when so many waterhole-dependent large grazers evolved in the Rift just before −2.35 myr, lightning strikes could result in booms for grazers and, secondarily, hominin meat-eaters.
3. *Why the Pleistocene climate swings had no effect on enlargement, once it began*: because mutations can quickly achieve fixation when aided by the feedback loops, the allele shifts are paced by the rate-limiter, the cosmic ray bombardment.
4. *Why enlargement was arguably linear*: there was a steady rate-limited accumulation of minor tweaks in favorable alleles for enlargement.
5. *When it ended*: within a mere 5 kyr at the end of a 2,300 kyr run, agriculture enlarged the core population to such an extent that the continuing meat booms became insignificant for further tweaking of core allele ratios.
6. *Why enlargement slowed in the regional side branches*: compared to the African Rift and the Red Sea coastal corridor, there were fewer effective settings (deep brush adjacent to grassland with grazers) for creating an effective feedback loop for the Neanderthals in forested Europe as well as in overly-grassy Asia, slowing enlargement in non-ancestral *Homo erectus*. Continuing immigration of Africans would still provide some regional enlargement trend.

Hitchhiking now seems promising to explore as an evolutionary path for syntax and the other structured intellectual functions: contingent planning, chains of logic, games with rules constraining moves, analogies that extend to parables, polyphonic music, and creativity’s eureka moments when incoherent mental assemblies suddenly all hang together, good to go.

Indeed, seeking coherence in a jumble of possibilities characterizes all the higher intellectual functions (Calvin 2004). While quick action may be needed in many situations, being unable to suspend judgment for long enough to explore less common possibilities often results in premature closure.

Changing traits continuously via selective survival is familiar from prey-predator arms races. But for exploring such evolutionary extravaganzas, where a novel capability seems to lack short-term incremental advantages, examining settings may be more productive. As sexual selection did for the extravagant peacock tail, so boom-and-bust feedback loops can surprise us with progressions that keep going automatically–as in that afore-mentioned neocortical preadaptation for the next new thing, in all the children who are above average.

## Acknowledgements

I thank my La Jolla colleagues for two decades of high-level discussions about human evolution, with long-term CARTA funding to UCSD/Salk from the G. Harold and Leila Y. Mathers Foundation. I thank my University of Washington colleagues James J. Anderson, Katherine Graubard, and Charles D. Laird for discussions about the manuscript, and my late colleague Arnold L. Towe for early discussions about grazer search images and how two-legged hunters escaped notice. Human and bonobo scaled photographs thanks to T. W. Deacon.

## Methods

The primary endocranial volume sources were the tables of Holloway et al (2004) and Shultz et al (2012) with some updating; the late australopithecine-sized skulls of *H. naledi* and *H. floresiensis* were intentionally omitted from Fig. 2.

## References

Aiello, L. C. & Wheeler, P. (1995) The expensive-tissue hypothesis: the brain and the digestive system in human and primate evolution. Current Anthropology 36:199–221, http://www.jstor.org/stable/2744104

Archer, W., et al (2014) Early Pleistocene aquatic resource use in the Turkana Basin. Journal of human evolution 77:74–87, doi: 10.1016/j.jhevol.2014.02.012

Bailey, G. N., Reynolds, S. C., King, G. C. P. (2010) Landscapes of human evolution: models and methods of tectonic geomorphology and the reconstruction of hominin landscapes. Journal of human evolution 60:257–280, doi:10.1016/j.jhevol.2010.01.004

BBC (2011) Three men and 15 lions. Video at http://www.bbc.co.uk/programmes/p00f0xy8

Beyin, A. (2013) A surface Middle Stone Age assemblage from the Red Sea coast of Eritrea: implications for Upper Pleistocene human dispersals out of Africa. Quaternary International 300:195–212, doi:10.1016/j.quaint.2013.02.015

Bloom, P. (2001) Précis of How Children Learn the Meanings of Words. Behavioral and Brain Sciences 24:1095–1103, doi: 10.1017/S0140525X01000139

Brace, C.L. (2000) Evolution in an anthropological view. Altamira Press. ISBN 0742502635

Bunn, H. T. (2001) Hunting, power scavenging, and butchering by Hadza foragers and by Plio-Pleistocene Homo. In: Meat-eating and human evolution. Stanford, C. B., & Bunn, H.T., eds. Oxford University Press, pp. 199–218. ISBN 9780195351293

Calvin, W.H. (1983) A stone’s throw and its launch window: timing precision and its implications for language and hominid brains. Journal of theoretical biology 104: 121–135, http://cogprints.org/21/1/1983JTheoretBiol.htm

Calvin, W.H. (2002) A brain for all seasons: human evolution and abrupt climate change. University of Chicago Press, at p.128. http://faculty.washington.edu/wcalvin/BrainForAllSeasons/

Calvin, W.H. (2004) A brief history of the mind. Oxford University Press. http://faculty.washington.edu/wcalvin/BHM

Calvin, W.H. (2017) Hitchhiking on the frontier: accelerating eusociality and other improbable evolutionary outcomes by trait hitchhiking in a boom-and-bust feedback loop. Preprint: bioarxiv, doi: 10.1101/053819

Calvin, W.H., Bickerton, D. (2000) Lingua ex machina. MIT Press, p.83, http://williamcalvin.org/LEM/

Carotenuto, F., et al (2016) Venturing out safely: The biogeography of Homo erectus dispersal out of Africa. Journal of Human Evolution, 95:1–12, doi: 10.1016/j.jhevol.2016.02.005

Cerling, T.E., et al (2015) Dietary changes of large herbivores in the Turkana Basin, Kenya from 4 to 1 Ma. Proceedings of the national academy of sciences USA 112:11467–11472, doi: 10.1073/pnas.13075112

Cerling, T.E., et al (2013) Stable isotope-based diet reconstructions of Turkana Basin hominins. Proceedings of the national academy of sciences USA 110:10501–10506, doi: 10.1073/pnas.1222568110

Conley, D., et al (2016) Assortative mating and differential fertility by phenotype and genotype across the 20th century. Proceedings of the national academy of sciences USA 113:6647–6652, doi: 10.1073/pnas.1523592113

Darwin, C. (1859) On the origin of species. 3rd edition. Murray.

de Ruiter, J., Weston, G. and Lyon, S. M. (2011) Dunbar’s number: group size and brain physiology in humans reexamined. American Anthropologist 113: 557–568, doi:10.1111/j.1548-1433.2011.01369.x

Dibble, H.L., et al. (2016) Major fallacies surrounding stone artifacts and assemblages. Journal of archaeological method and theory 1-39, doi:10.1007/s10816-016-9297-8

Dohrn, A. (1875) Der ursprung der wirbelthiere und das princip des functionswechsels: genealogische skizzen. Leipzig: Engelmann.

Dunbar, R. (1996) Grooming, gossip and the evolution of language. Harvard University Press.

Dunbar, R.I.M. (2004) Gossip in evolutionary perspective. Review of general psychology 8: 100–110, doi: 10.1037/1089-2680.8.2.100.

Falk, D. (1990) Brain evolution in Homo: the “radiator” theory. Behavioral and Brain Sciences 13:333–344.

Faure, H., Walter, R.C., Grant, D.R. (2002) The coastal oasis: ice age springs on emerged continental shelves. Global and planetary change 33:47–56, doi: 10.1016/S0921-8181(02)00060-7

Fenner, J.N. (2005) Cross-cultural estimation of the human generation interval for use in genetics-based population divergence studies. American Journal of Physical Anthropology 125:415–423, doi: 10.1002/ajpa.20188

Ferraro, J.V., et al. (2013) Earliest archaeological evidence of persistent hominin carnivory. PLoS ONE 8:e62174, doi:10.1371/journal.pone.

Finlay, B.L. (2004) The Calvinist cortex: penetrating evolutionary predestination. Cortex 40:577–579.

Finlay, B.L., & Darlington, R.B. (1995) Linked regularities in the development and evolution of mammalian brains. Science 268:1578–1584, doi: 10.1126/science.7777856

Finlay, B.L., & Uchiyama, R. (2015) Developmental mechanisms channeling cortical evolution. Trends in neuroscience 36:69–76, doi: 10.1016/j.tins.2014.11.004

Glasser, M.F., et al (2016) A multi-modal parcellation of human cerebral cortex. Nature 536:171–178, doi: 10.1038/nature18933M3

Harmand, S., et al (2015) 3.3-million-year-old stone tools from Lomekwi 3, West Turkana, Kenya. Nature 521:310–315, doi: 10.1038/nature14464

Heape, W. (1899) Note on the fertility of different breeds of sheep, with remarks on the prevalence of abortion and barrenness therein. Proceedings of the Royal Society London 65, 99–111

Higham, T., et al (2014) The timing and spatiotemporal patterning of Neanderthal disappearance. Nature, 512:306–309, doi: 10.1038/nature13621.

Holloway, R.L., Yuan, M.S., Broadfield, D.C. (2004) The human fossil record: brain endocasts - the paleo-neurological evidence, volume three. New York: Wiley, at Appendix 1, pp. 295.

Kong, A., et al. (2012). Rate of de novo mutations and the importance of father’s age to disease risk. Nature 488, pp. 471–475, doi: 10.1038/nature11396.

Kuhl, P.K. (2004) Early language acquisition: cracking the speech code. Nature reviews neuroscience, 5, pp. 831–843, doi:10.1038/nrn1533

Lieberman, D.E. (1998) Sphenoid shortening and the evolution of modern human cranial shape. Nature 393:158–162, doi:10.1038/30227

Lieberman, D.E., Bramble, D.M., Raichlen, R.A., Shea, J.J. (2009) Brains, brawn, and the evolution of human endurance running capabilities. In: Grine, F.E., Fleagle, J.G., Leakey, R.E., editors. The first humans – origin and early evolution of the genus Homo. Springer Netherlands; pp. 77–92. Link: link.springer.com/chapter/10.1007%2F978-1-4020-9980-9_8.

Lisiecki, L.E., Raymo, M.E. (2005) A Pliocene-Pleistocene stack of 57 globally distributed benthic δ18O records. Paleoceanography 20: PA1003. doi: 10.1029/2004PA001071. Fig. 2B is replotted from their database.

Lund, J. S., Boothe, R. G. and Lund, R. D. (1977) Development of neurons in the visual cortex (area 17) of the monkey (Macaca nemestrina): A Golgi study from fetal day 127 to postnatal maturity. Journal of Comparitive Neurology, 176: 149–187, doi:10.1002/cne.901760203

Mackey, W.E., Winawer, J., Curtis, C.E. (2017) Visual field map clusters in human frontoparietal cortex. Preprint at http://biorxiv.org/content/biorxiv/early/2016/10/27/083493.full.pdf

Marlowe, F.W. (2005) Hunter-gatherers and human evolution. Evolutionary anthropology 14:54–67, doi: 10.1002/evan.20046

McDougall, I., Brown, F.H., & Fleagle, J.G. (2008) Sapropels and the age of hominins Omo I and II, Kibish, Ethiopia. Journal of human evolution 55:409–420, doi: 10.1016/j.jhevol.2008.05.012

Messager, E., Lordkipanidze, D., Kvavadze, E., Ferring, C.R., Voinchet, P. (2010) Palaeoenvironmental reconstruction of Dmanisi site (Georgia) based on palaeobotanical data. Quaternary International 223:20–2, doi: 10.1016/j.quaint.2009.12.016.

Moorjani, P., Gao, Z., Przeworski, M. (2016) Human germline mutation and the erratic evolutionary clock. PLoS Biol 14:e2000744, doi: 10.1371/journal.pbio.2000744

Morgan, E. (1982) The aquatic ape. Stein & Day, p.69.

Orban, G.A., Fausto, C. (2014) The neural basis of human tool use. Frontiers in psychology 5:1–12, doi: 10.3389/fpsyg.2014.00310

Pääbo, S. (2014) Neanderthal man: in search of lost genomes. Basic Books.

Pobiner, B. (2016) Meat-eating among the earliest humans. American Scientist 104:110–116, doi: 10.1511/2016.119.110

Prat, S. (2016) First hominin settlements out of Africa. Tempo and dispersal mode: Review and perspectives. Comptes Rendus Palevol, doi: 10.1016/j.crpv.2016.04.009

Roach, N.T., Venkadesan, M., Rainbow, M.J., Lieberman, D.E. (2013) Elastic energy storage in the shoulder and the evolution of high-speed throwing in Homo. Nature 498: 483–486, doi:10.1038/nature12267.

Rosenberg, K. R. (1992) The evolution of modern human childbirth. American journal of physical anthropology 35:89–124, doi:10.1002/ajpa.1330350605

Ruff, C. B. (2006) Gracilization of the modern human skeleton. American Scientist 94:508–514, doi: 10.1511/2006.62.1007

Schoe, M., De Iongh, H. H. and Croes, B. M. (2009) Humans displacing lions and stealing their food in Bénoué National Park, North Cameroon. African journal of ecology, 47: 445–447. doi:10.1111/j.1365-2028.2008.00975.x

Schoeninger, M.J. (2012) Palaeoanthropology: The ancestral dinner table. Nature 487:42–43, doi:10.1038/487042a

Seymour, R. S., Bosiocic, V., Snelling, E. P. (2016) Fossil skulls reveal that blood flow rate to the brain increased faster than brain volume during human evolution. Royal Society Open Science 3:160305, doi:10.1098/rsos.160305

Shipman, P. (1986) Scavenging or hunting in early hominids: theoretical framework and tests. American Anthropologist 88:27–43, doi: 10.1525/aa.1986.88.1.02a00020

Shipman, P., Walker, A. (1989) The costs of becoming a predator. Journal of human evolution 18:373–392

Shultz, S., Nelson, E., Dunbar, R.I.M (2012) Hominin cognitive evolution: identifying patterns and processes in the fossil and archaeological record. Philosophical Transactions of the Royal Society B: Biological Sciences 367: 2130–2140, doi: 10.1098/rstb.2012.0115

Teffer, K., Semendeferi, K. (2012) Human prefrontal cortex: evolution, development, and pathology. In: Hofman M.A., Falk D., editors. Evolution of the primate brain: from neuron to behavior. London: Elsevier; pp. 191–218.

Trinkel, M., Kastberger, G. (2005) Competitive interactions between spotted hyenas and lions in the Etosha National Park, Namibia. African Journal of Ecology 43:220–224. doi: 10.1111/j.1365-2028.2005.00574.x.

Turchin, P. (2015) Ultrasociety: How 10,000 years of war made humans the greatest cooperators on earth. Beresta Books.

van der Made, J. (2011) Biogeography and climatic change as a context to human dispersal Out of Africa and within Eurasia. Quaternary Science Reviews 30:1353–1367, doi: 10.1016/j.quascirev.2010.02.028

Villmoare, B., et al (2015) Early Homo at 2.8 Ma from Ledi-Geraru, Afar, Ethiopia. Science 347:1352–1355, doi: 10.1126/science.aaa1343

Walter, R. C., et al (2000) Early human occupation of the Red Sea coast of Eritrea during the last interglacial. Nature 405:65–69, doi:10.1038/35011048

Wang, L., et al (2015) Probabilistic maps of visual topography in human cortex. Cerebral Cortex 25:3911–3931, doi:10.1093/cercor/bhu277

Witelson, S.F., Beresh, H., Kigar, D.L. (2006) Intelligence and brain size in 100 postmortem brains: sex, lateralization and age factors. Brain 129:386–398, doi: 10.1093/brain/awh696.

